# Heterotypic vaccination responses against SARS-CoV-2 Omicron BA.2

**DOI:** 10.1101/2022.03.22.485418

**Authors:** Zhenhao Fang, Lei Peng, Qianqian Lin, Liqun Zhou, Luojia Yang, Yanzhi Feng, Ping Ren, Paul A. Renauer, Jonathan J. Park, Xiaoyu Zhou, Craig B. Wilen, Sidi Chen

**Author notes:** Co-first authors (Zhenhao Fang, Lei Peng). Correspondence: SC, (lab).

## Abstract

The Omicron sub-lineage BA.2 of SARS-CoV-2 has recently become dominant across many areas in the world in the on-going waves of COVID-19. Compared to the ancestral/wild-type (WT) virus, Omicron lineage variants, both BA.1 and BA.2, contain high number of mutations, especially in the spike protein, causing significant immune escape that leads to substantial reduction of vaccine and antibody efficacy. Because of this antigenic drift, BA.2 exhibited differential resistance profile to monoclonal antibodies than BA.1. Thus, it is important to understand whether the immunity elicited by currently available vaccines are effective against the BA.2 subvariant. We directly tested the heterotypic vaccination responses against Omicron BA.2, using vaccinated serum from animals receiving WT- and variant-specific mRNA vaccine in lipid nanoparticle (LNP) formulations. Omicron BA.1 and BA.2 antigen showed similar reactivity to serum antibodies elicited by two doses of WT, B.1.351 and B.1.617 LNP-mRNAs. Neutralizing antibody titers of B.1.351 and B.1.617 LNP-mRNA were ~2-fold higher than that of WT LNP-mRNA. Both homologous boosting with WT LNP-mRNA and heterologous boosting with BA.1 LNP-mRNA substantially increased waning immunity of WT vaccinated mice against both BA.1 and BA.2 subvariants. The BA.1 LNP-mRNA booster was ~3-fold more efficient than WT LNP-mRNA at elevating neutralizing antibody titers of BA.2. Together, these data provided a direct preclinical evaluation of WT and variant-specific LNP-mRNAs in standard two-dose and as boosters against BA.1 and BA.2 subvariants.

---

Coronavirus disease 2019 (COVID-19) pandemic has taken away over 6 million lives in the past two years, and continues to pose a significant threat to the world due to the increased transmissibility, infectivity and immune evasion of continuously emerging variants of severe acute respiratory syndrome coronavirus 2 (SARS-CoV-2)^1^. Within weeks since its first identification in southern Africa, the newly emerged variant of concern, Omicron (B.1.1.529) has become the dominant variant and spread rapidly worldwide^2^. The spread of Omicron initial form BA.1 was followed by a rapid rise of an Omicron sub-lineage BA.2, which is now also designated as a variant of concern (VoC)^3^ and has represented more than 70% North America cases and 80% global cases^2^, eclipsing the once-dominant BA.1. The on-going “fifth wave” and “sixth wave” of COVID-19 are predominantly associated with BA.2 and have claimed hundreds of thousands of lives to date, especially in Asia and Europe at the time of this study^4,5^.

Compared to the ancestral / wild-type (WT) virus, Omicron variants contain an alarming number of mutations (over 30 mutations) in spike protein, which is the primary target of clinical antibodies and vaccines. The substantial differences between WT and Omicron spike lead to extensive immune escape of Omicron from WT mRNA vaccine^6^, which prompt the idea of developing Omicron-specific vaccines. We generated several COVID variant-specific mRNA vaccine candidates (including BA.1 subvariant)^7,8^ which were designed based on variants’ spike stabilized by six proline mutations^9^. Variant-specific vaccine candidates, or lipid nanoparticle (LNP)- mRNAs, unequivocally exhibited varying degrees of advantage over WT LNP-mRNA in terms of eliciting neutralizing antibody against cognate variant antigens^7,8^. Moreover, immune profiling of Omicron BA.1 LNP-mRNA showed a significant boosting effect on waning immune response of WT LNP-mRNA vaccinated mice to both Delta and Omicron BA.1 variants.

BA.1 and BA.2 subvariants share 21 mutations, but differ in 25 sites (**Fig. 1a-1b**). Because of this antigenic drift, BA.2 exhibited differential resistance profile to monoclonal antibodies than BA.1^10^. The significant difference of BA.1 and BA.2 spikes raises a number of profound questions. For instance, how potent is the immunity elicited by heterotypic vaccination, with WT or variant specific LNP-mRNAs, against BA.2 subvariant? How does this immune response compare to the response to BA.1? Does heterologous boosting with WT plus BA.1 LNP-mRNA or homologous boosting with WT LNP-mRNA remain effective against BA.2?

**Figure 1.**
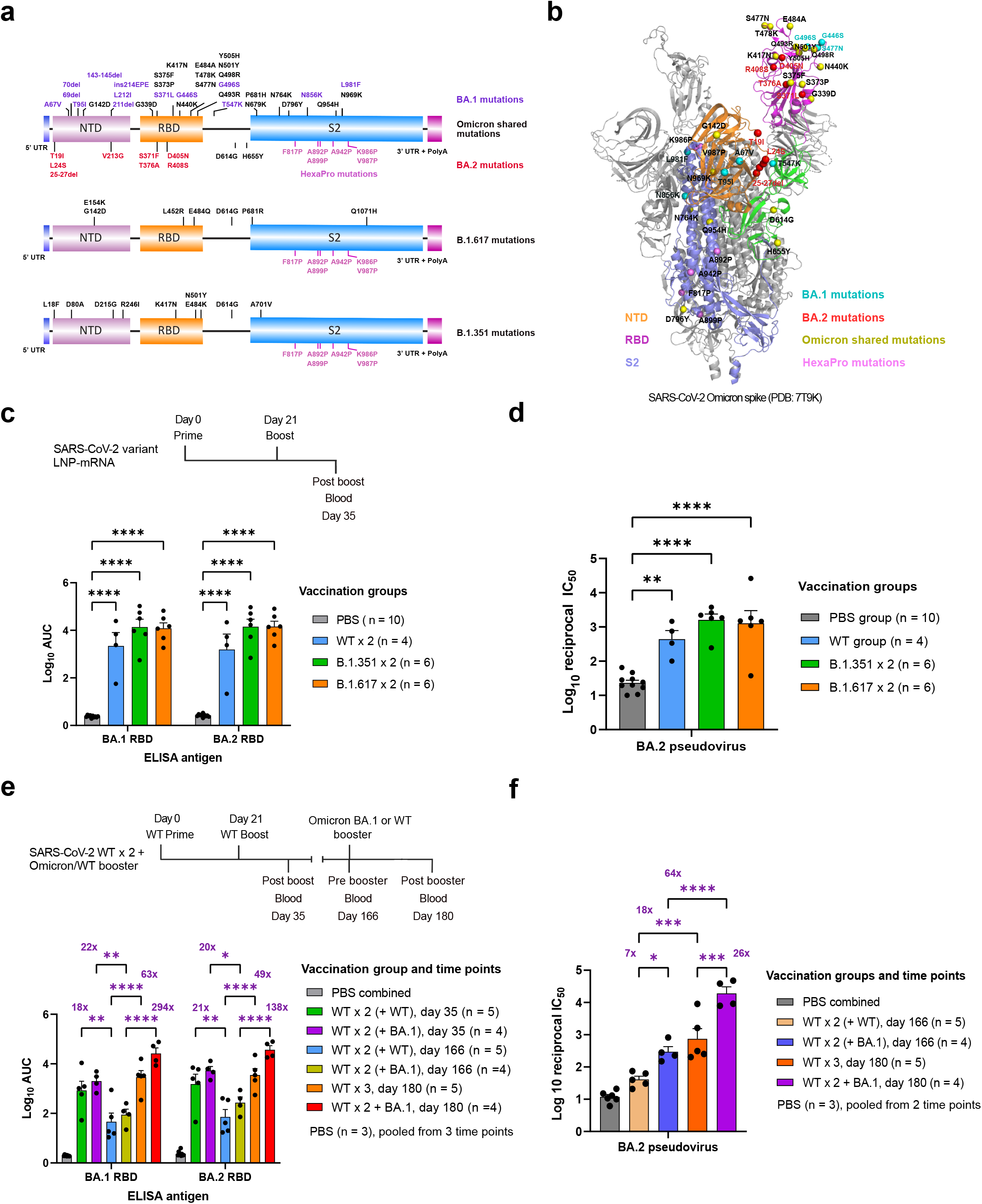
WT and variant-specific LNP-mRNA elicited potent antibody response against Omicron BA.1 and BA.2 sublineages. **a**, Schematics showing variant mutation distribution on spike sequences used in the variant specific vaccine design. **b**, Omicron BA.1 and BA.2 mutations were displayed in one protomer of Omicron BA.1 spike trimer (PDB: 7T9K). **c**, Comparison of antibody response induced by two doses of 1μg WT, B.1.351 or B.1.617 LNP-mRNA at 21 days interval. Vaccination scheme and blood collection time were shown on the time axis (top). Antibody titers were determined by area under curve (AUC) of ELISA titration curves in Figure S1. The number of animals in each vaccination group were shown as n in the bracket. **d**, Neutralization of Omicron BA.2 pseudovirus by serum samples from mice vaccinated with 1μg WT, B.1.351 or B.1.617 LNP-mRNA as illustrated in **Fig. 1c**. The neutralizing titers were quantified by log_10_ reciprocal IC50 and calculated from titrations in **Fig. S2**. **e**, BA.1 and WT boosters strengthened waning immunity against both Omicron BA.1 and BA.2 variants. Vaccination scheme and blood collection time were shown on the time axis (top). ELISA antibody titers of samples from mice sequentially vaccinated with two doses of 1μg WT LNP-mRNA followed by 10μg WT (WT × 3, n = 5) or Omicron BA.1 (WT × 2 + BA.1, n = 4) LNP-mRNA boosters. The pre-booster groups (day 35 and day 166) to receive WT or BA.1 boosters were denoted as WT × 2 (+ WT) and WT × 2 (+ BA.1) respectively. **f**, Neutralization of Omicron BA.2 pseudovirus by plasma samples from mice before and after receiving WT or Omicron BA.1 LNP-mRNA boosters as illustrated in **Fig. 1e**. Individual data points represent value from each mouse sample and are shown on dot-bar plots as mean ± s.e.m.. Data points of PBS group showed no statistical difference between collection time points and were combined to one group in graph EF. To assess statistical significance, two-way ANOVA with Tukey's multiple comparisons test was used. Statistical significance labels: * p < 0.05; ** p < 0.01; *** p < 0.001; **** p < 0.0001. Non significant comparisons are not shown. Only comparisons between adjacent time points or groups of same time point were shown in Fig. 1e-1f.

To answer these questions, we first characterized the antibody response induced by WT or variant specific LNP-mRNAs to Omicron BA.2 sublineage and compared it with immune response to BA.1. Samples used for BA.2 characterization were collected from mice that received two doses of 1μg WT, B.1.351 or B.1.617 LNP-mRNAs ^8^. All three LNP-mRNA including WT, B.1.351 and B.1.617 elicited significant antibody response to BA.2 (**Fig. 1c-1d**). Both B.1.351 and B.1.617 LNP-mRNA treatment group showed a trend of higher binding and neutralizing titers than WT group, albeit insignificant. Because of selection pressure, emerging variants often retain some signature mutations conferring fitness advantage from past variants^11^. BA.2 shares 3 mutations with B.1.351 (K417N, N501Y, D614G) and B.1.617 (G142D, D614G, P681R), which may explain why the antibody response to BA.2 was higher in these two variants LNP-mRNA groups compared to WT LNP-mRNA (**Fig. 1a**). In all three vaccination groups, antibody response to BA.2 was similar to that of BA.1 (**Fig. 1c**), suggesting approximately equal reactivity of BA.1 and BA.2 to heterotypic vaccination by WT, B.1.351 and B.1.617 LNP-mRNA. It is worth noting that both BA.1 and BA.2 share the same 3 mutations with B.1.351 and B.1.617, which contributed to the conserved cross reactivity of variant LNP-mRNA to two Omicron sublineages.

Given the BA.2 neutralization titer advantage of variant LNP-mRNA over WT counterpart, we went on to profile the antibody response of BA.1 LNP-mRNA to BA.2 subvariant. To model the real-world scenario of boosting waning immunity of general population receiving WT mRNA vaccines^12,13^, we sought to investigate the effect of homologous boosting with WT LNP-mRNA or heterologous boosting with BA.1 LNP-mRNA on waning immunity of WT vaccinated animals against Omicron BA.2. The overall antibody titer changes over time in matched booster groups showed similar trend within BA.1 and BA.2 ELISA datasets (**Fig. 1e**). A 20-fold time-dependent decrease in antibody titer was observed over 4 months (day 35 vs. day 166) in both BA.1 and BA.2 datasets, suggesting evident and comparable waning immunity for the two Omicron sublineages. When comparing the boosting effect of WT and BA.1 LNP-mRNA, BA.1 LNP-mRNA consistently showed a better performance than WT in BA.1 and BA.2 datasets. The antibody titer increases by BA.1 LNP-mRNA were 293-fold (fold change = titers ratio - 1) and 137-fold for BA.1 and BA.2 antigens respectively, while the ones mediated by WT LNP-mRNA were 62-fold and 48-fold. Comparing to BA.1 antigen, both WT and BA.1 LNP-mRNA showed weaker boosting effects on BA.2 antigen and this effect reduction was more apparent for BA.1 LNP-mRNA than WT LNP-mRNA. As the post-booster titers against BA.1 and BA.2 were quite similar, this reduction was mainly due to higher pre-booster titers against BA.2 antigen, although such pre-booster titer difference between BA.1 and BA.2 did not reach statistical significance. The data from pseudovirus neutralization assay of BA.2 correlated well with corresponding ELISA data and strengthened the forementioned findings in ELISA (**Fig. S3**). The neutralizing titer enhancement mediated by WT and BA.1 boosters were 18-fold (p < 0.001) and 63-fold (p < 0.0001), respectively (**Fig. 1f**). Importantly, the heterotypic vaccination by Omicron BA.1 LNP-mRNA vaccine booster is more efficient at boosting neutralizing titers than WT LNP-mRNA booster (comparing boosting effect of WT vs. BA.1, 64/18=3.6, **Fig. 1f**). These data highlight the benefit of receiving booster shots and advantage of BA.1-specific vaccine over WT vaccine against BA.2.

In summary, our data showed a significant drop of antibody titers over time and clear benefit of heterotypic vaccination by WT and BA.1 LNP-mRNA boosters on both BA.1 and BA.2 subvariants, which justify and necessitate the use of homologous WT or heterologous BA.1 boosters in order to curb the fast spread of Omicron subvariants. The heterologous booster by BA.1 vaccination on top of the two-dose WT vaccination may provide stronger benefit against the BA.2 variant, which is the current global dominant VoC. The remarkable antigenic drift of emerging variants from WT virus renders many existing clinical antibodies and vaccines suffer from efficacy loss, which is especially evident for Omicron BA.1 and BA.2 subvariants. To prevent this ever-evolving enemy breaking through our line of defense, we generated and characterized a number of variant-specific LNP-mRNAs, including B.1.351, B.1.617 and BA.1. Because of shared mutations with BA.1 or BA.2 sublineages, these variant-specific LNP-mRNA displayed better performance of inducing neutralizing antibodies than WT LNP-mRNA in booster and non-booster settings. Rapid development and preclinical characterization of these variant-specific LNP-mRNAs would benefit the development of mRNA vaccines targeting the evolving variants.

## Supplemental figure legend

**Figure S1.**
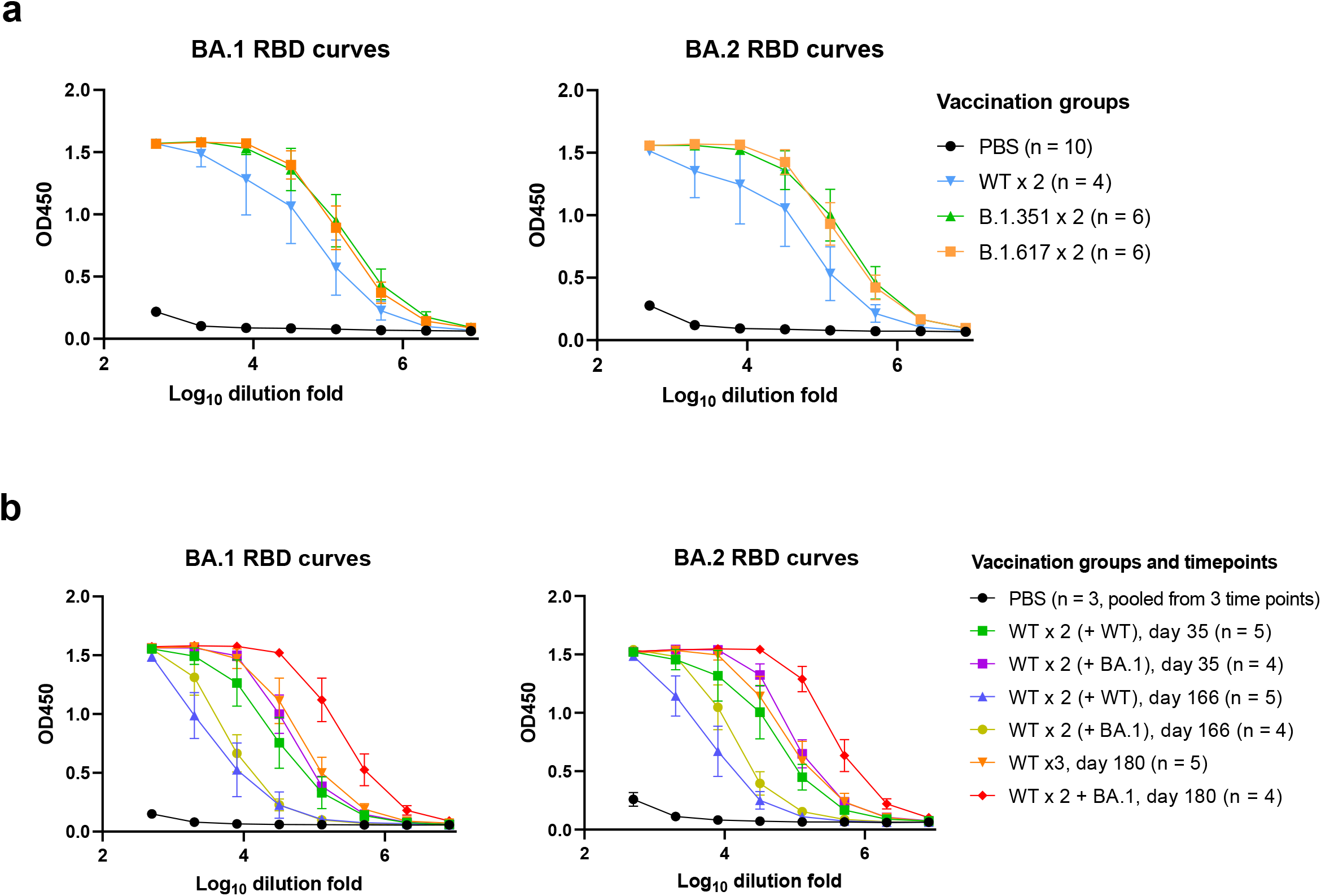
ELISA dose-response curves of serially diluted plasma or sera collected at indicated time points from mice vaccinated with WT or variant specific LNP-mRNA. **a**, Titration curves against BA.1 (left) and BA.2 (right) RBDs by samples from mice immunized with two doses of 1ug WT, B.1.351 or B.1.617 LNP-mRNAs. **b**, Titration curves against BA.1 (left) and BA.2 (right) RBDs by mice samples before and after 10ug WT or BA.1 LNP-mRNA booster shots. The average OD450 response were shown as mean ± s.e.m. and plotted against serial log_10_-transformed sample dilution points.

**Figure S2.**
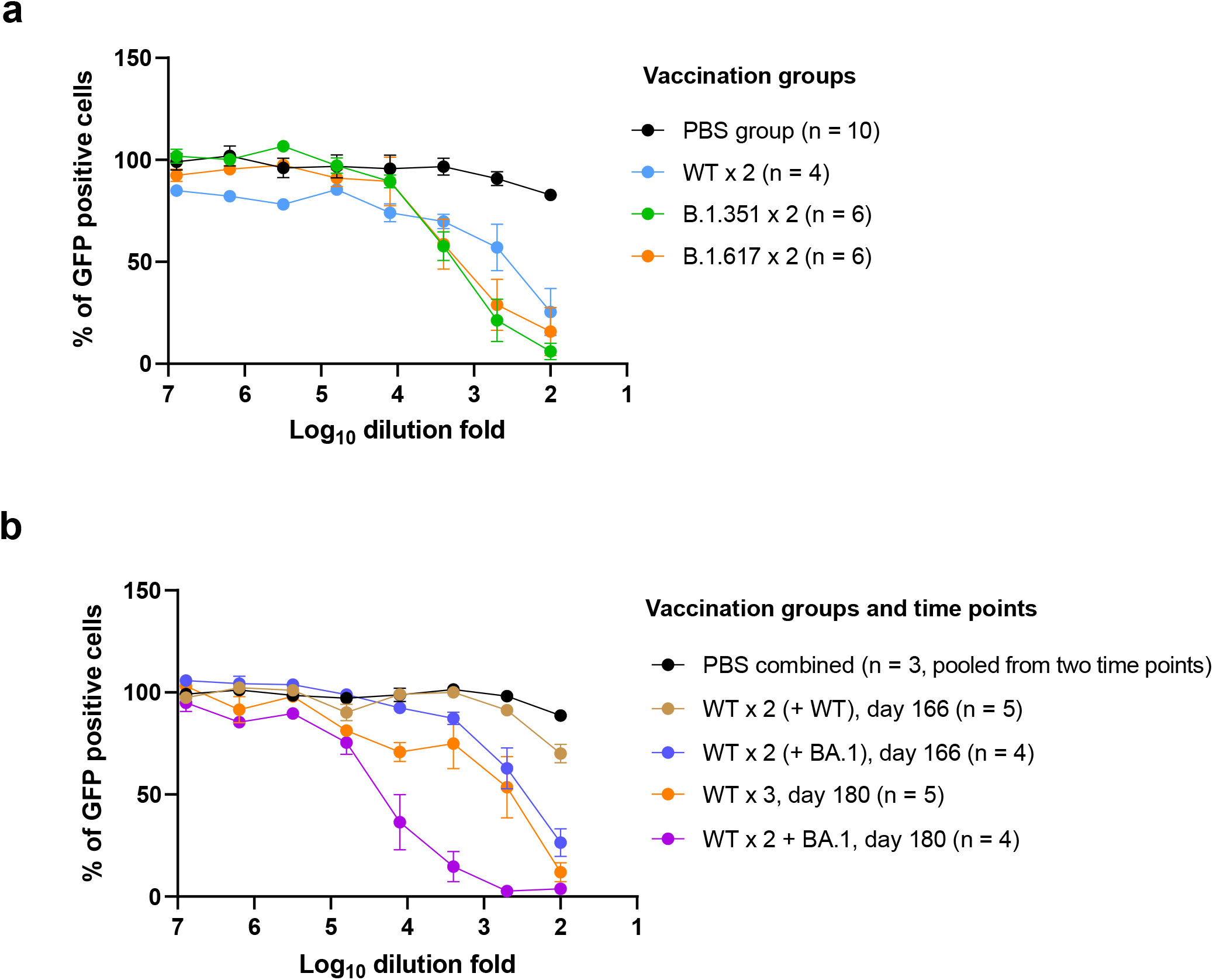
Neutralization titration curves of serially diluted plasma or sera collected at indicated time points from mice vaccinated with WT or variant specific LNP-mRNA. **a**, Neutralization curves of BA.2 pseudovirus by samples from mice immunized with two doses of 1ug WT, B.1.351 or B.1.617 LNP-mRNAs. **b**, Neutralization curves of BA.2 pseudovirus by samples before and after 10ug WT or BA.1 LNP-mRNA booster shots. The average GFP positive rates or pseudovirus infection rates were shown as mean ± s.e.m. and plotted against serial log10-transformed sample dilution points.

**Figure S3.**
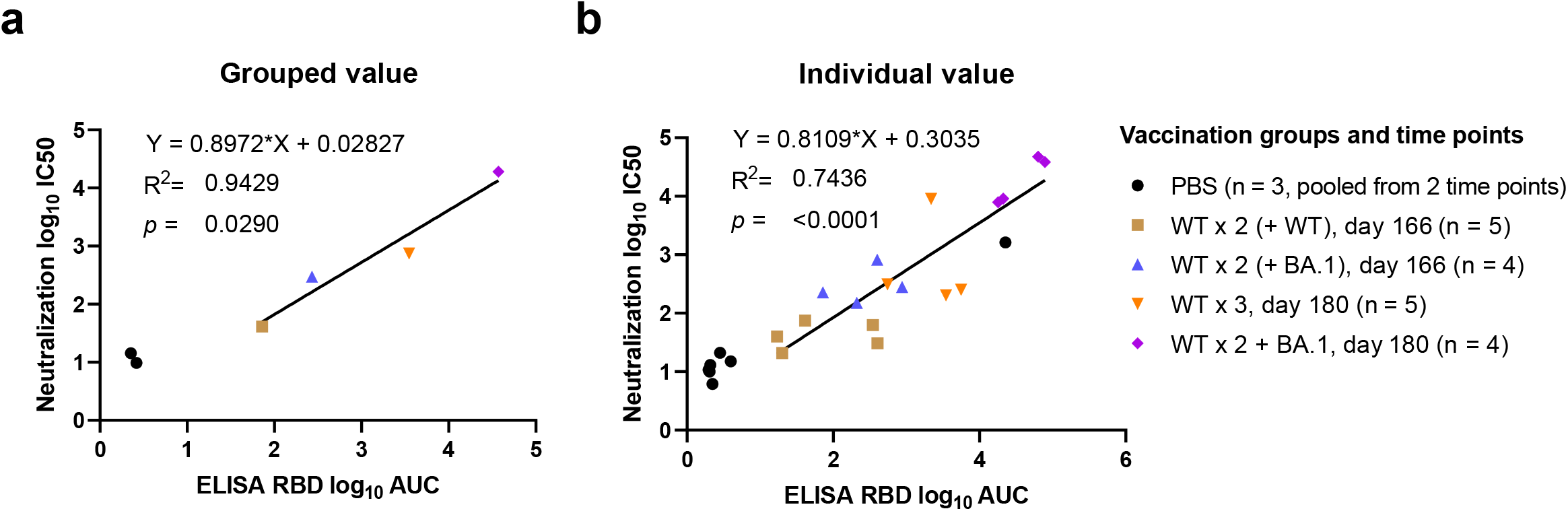
Correlation between antibody titers measured by ELISA and pseudovirus neutralization assay. Pseudovirus neutralizing antibody titers were shown on y axis as log10 reciprocal IC50 and plotted against ELISA binding antibody titers on x axis (log_10_ AUC). Titer values were either from mean of matched vaccination group (a) or individual animal (b).

**Figure S4.**
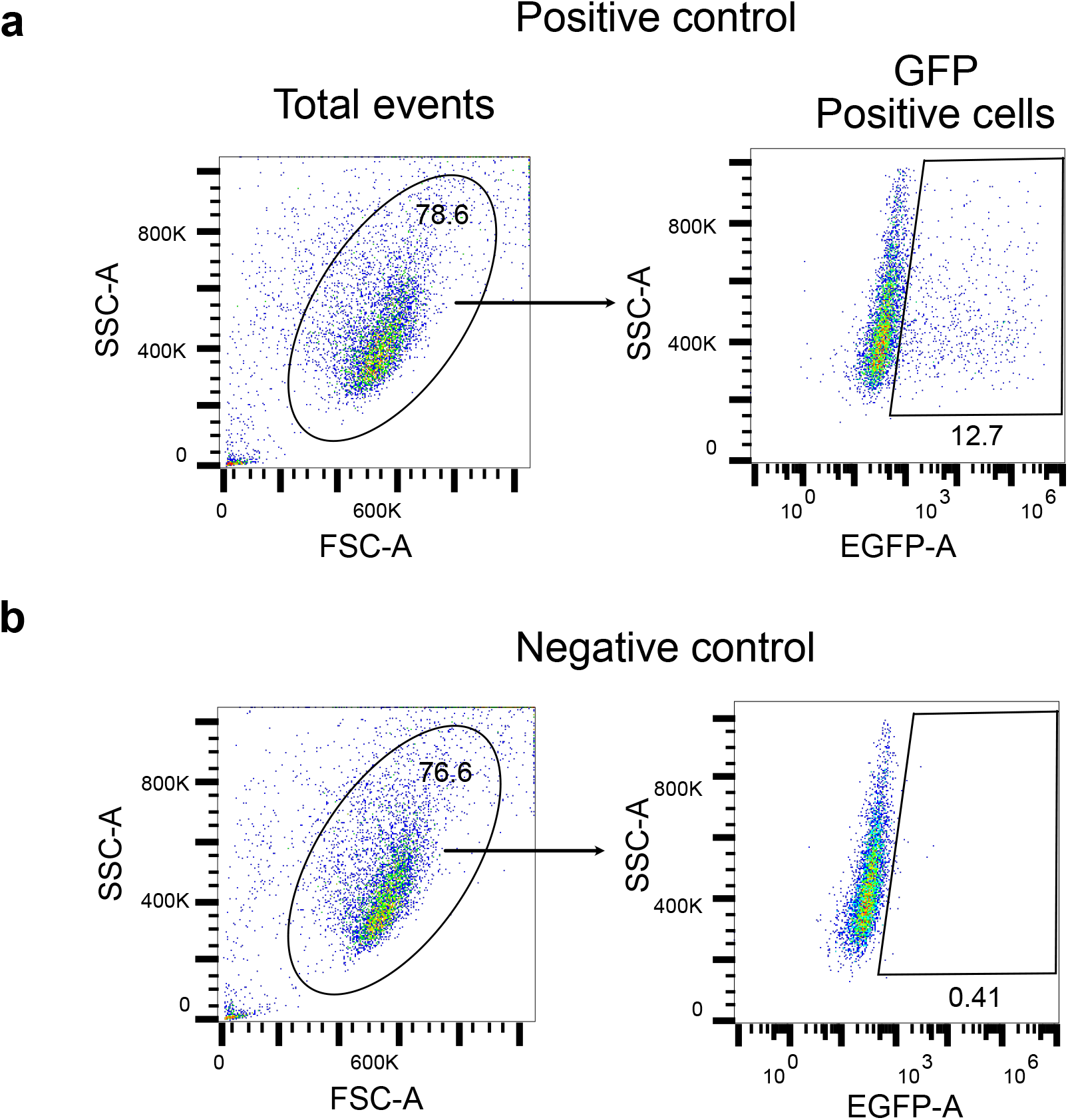
Representative flow cytometry gating strategy used in pseudovirus neuralization assay for detecting GFP positive or infected cells.

## Methods

### Molecular cloning and mRNA transcription

The coding sequence of Omicron BA.2 spike were derived from isolates in GISAID EpiCoV database (EPI_ISL_6795834.2). The spike plasmids were linearized by restriction enzymes and transcribed to mRNA by in vitro T7 RNA polymerase (NEB, Cat # E2060S) as previously described^7,8^.

### Cell culture

293T and hACE2-293FT cells were maintained in Dulbecco’s minimal essential medium (DMEM, Fisher) supplemented with 10% fetal bovine serum (Hyclone) and penicillin (100 U/ml)-streptomycin (100 ug/ml). Cells were split ever other day at a 1:4 ratio when confluency is over 90%.

### Lipid nanoparticle mRNA preparation

The lipid nanoparticle mRNA were prepared as previously described^7,8^. In brief, lipid mixture was dissolved in ethanol and mixed with mRNA in pH 5.2 sodium acetate. The mRNA encapsulated by LNP (LNP-mRNA) was then exchanged to PBS using 100kDa Amicon filter (Macrosep Centrifugal Devices 100K, 89131-992). The DLS device was used to validate the size distribution of LNP-mRNA (DynaPro NanoStar, Wyatt, WDPN-06). The encapsulation rate and mRNA amount were determined by Quant-iT™ RiboGreen™ RNA Assay (Thermo Fisher).

### Animal vaccination

Animal immunization were performed previously on 6-8 weeks female C57BL/6Ncr mice purchased from Charles River in two sets of experiments: 1) sequential vaccination with two doses of 1μg WT LNP-mRNA followed by 10μg Omicron BA.1 or WT boosters^7^; 2) vaccination with two doses of 1μg WT, B.1.351, B.1.617 LNP-mRNA^8^. Retro-orbital blood were collected two weeks post boost (2^nd^ dose, day 35), right before boosters (day 166), and two weeks post boosters (3^rd^ dose, day 180).

### ELISA and Neutralization assay

The binding and neutralizing antibody titers were determined by ELISA and pseudovirus neutralization assay as previously described^7,8^. The Omicron BA.1 RBD and BA.2 RBD used in ELISA were purchased from Sino Biological (Cat. No. 40592-V08H121) and AcroBiosystems (Cat. No. SPD-C522g-100ug) respectively. The pseudovirus plasmids were generated based on the WT plasmid which was a gift from Dr. Bieniasz’s lab^14^.

## Data availability

All source data and statistics are provided in this article and its supplementary table excel file. Additional information related to this study are available from the corresponding author upon reasonable request.

## Code availability

No custom code was used in this study.

## Acknowledgements

We thank various members from our labs for discussions and support. We thank Drs. M. Müschen, and L. Chen for generously providing equipment support. We thank staffs from various Yale core facilities (Keck, YCGA, HPC, YARC, CBDS and others) for technical support. We thank various support from Department of Genetics; Institutes of Systems Biology and Cancer Biology; Dean’s Office of Yale School of Medicine and the Office of Vice Provost for Research.

## Author Contributions

ZF: design of the study groups, constructs, cloning, vaccine system development, ELISA, data analysis, figure prep, and writing

LP: vaccination system development, immunization, sample collection, neutralization, and data analysis

QL, LZ, LY, YF, PR: assisting experiments, resources

PAR, JJP, XZ: vaccination system development

CBW: resources, supervision

SC: conceptualization, overall design, funding, supervision

## Notes

### Competing Interest Statement

The authors have declared no competing interest.

